# CA1 20-40 Hz oscillatory dynamics reflect trial-specific information processing supporting nonspatial sequence memory

**DOI:** 10.1101/2020.03.10.985093

**Authors:** Sandra Gattas, Gabriel A. Elias, Michael A. Yassa, Norbert J. Fortin

## Abstract

The hippocampus is known to play a critical role in processing information about temporal context. However, it remains unclear how hippocampal oscillations are involved, and how their functional organization is influenced by connectivity gradients. We examined local field potential activity in CA1 as rats performed a complex odor sequence memory task. We find that odor sequence processing epochs were characterized by increased power in the 4-8 Hz and 20-40 Hz range, with 20-40 Hz oscillations showing a power gradient increasing toward proximal CA1. Running epochs were characterized by increased power in the 8-12 Hz range and across higher frequency ranges (>24 Hz), with power gradients increasing toward proximal and distal CA1, respectively. Importantly, 20-40 Hz power increased with knowledge of the sequence and carried trial-type-specific information. These results suggest that 20-40 Hz oscillations are associated with trial-specific processing of nonspatial information critical for order memory judgments.

## Introduction

Brain oscillations are associated with many cognitive functions (Buzsaki et al., 2013; Colgin, 2016; Pesaran et al., 2018) and are thought to reflect complex interactions of neural activity from diverse populations of interconnected neurons (Pesaran et al., 2018). For instance, it is well established that the hippocampus is critical for spatial learning and memory (O’Keefe & Nadel, 1978), and that distinct oscillatory states are observed in hippocampal subregion CA1 during spatial navigation in rodents (see Colgin, 2016). In addition to the prominent theta rhythm (8-12 Hz), there is also evidence that the CA1 network exhibits transient increases in slow gamma (25-55 Hz) and fast gamma (60-100 Hz) power during running. In fact, a landmark study by Colgin and colleagues (2009) showed that CA1’s slow gamma oscillations are coherent with CA3 activity, whereas CA1’s fast gamma oscillations are coherent with entorhinal activity, suggesting these brief oscillatory states reflect retrieval and encoding processes, respectively. However, establishing a direct link between these oscillatory states and specific forms of information processing remains challenging, as such paradigms tend to have poor control over the timing at which information is encoded or retrieved.

In addition to spatial information, accumulating evidence indicates that the hippocampus plays a key role in the processing of temporal information. Consistent with its unique architecture and connectivity (McNaughton and Morris, 1987; Lisman 1999; Foster and Knierim 2012; Buzsáki and Tingley 2018), a growing literature shows the hippocampus is critical for remembering sequences of nonspatial events (Fortin et al. 2002; Kesner et al. 2002; Allen and Fortin, 2013; Eichenbaum, 2014) and that hippocampal neurons code for temporal relationships among such events (MacDonald et al., 2011; Allen et al. 2016; Shahbaba et al., 2019). However, little is known about the oscillatory dynamics associated with this fundamental type of information processing in the hippocampus. For instance, previous studies have shown theta oscillations in CA1 while rodents sampled task-relevant nonspatial stimuli, though the frequency range tends to be lower than during running (∼4-8 Hz in tasks using olfactory stimuli; Martin et al., 2007; Igarashi et al., 2014; Allen et al., 2016). In addition, there is evidence that the hippocampus exhibits oscillations in the beta range (∼20-40Hz) during the processing of odor information (Martin et al., 2007; Igarashi et al., 2014; Allen et al., 2016), and that this signal varies across the proximodistal axis of CA1 (higher power in distal than proximal CA1; Igarashi et al., 2014). However, it remains unclear whether these oscillations are associated with specific cognitive processes or aspects of performance, and whether proximodistal gradients in oscillatory power reflect the modality of the stimulus or vary with task demands. Further, it remains to be determined whether oscillations observed during spatial exploration extend to the processing of nonspatial information.

To address these important issues, we examined local field potential (LFP) activity in CA1 as rats performed a hippocampus-dependent odor sequence memory task (Fig 1). Importantly, this complex task offers precise time-locking to stimulus presentations and responses, as well as distinct trial types, contrasts and time windows associated with distinct cognitive demands. As in our previous work (Allen et al., 2016), we report prominent oscillations in the 20-40 Hz and 4-8 Hz frequency ranges during the odor sequence processing periods. Here we extend these results by demonstrating that the same electrodes exhibited a distinct spectral content in a different state (running on the track), which was characterized by high power in the 8-12 Hz band and a broad but modest increase in power for frequencies above 24 Hz. In both behavioral states, the power of recruited oscillations was found to vary along the proximodistal axis of CA1. We also made two additional contributions to our understanding of 20-40 Hz oscillations to hippocampal function. First, we discovered that 20-40 Hz oscillations were linked with sequence memory performance, whereas oscillations in other frequency ranges did not show a significant association. 20-40 Hz power increased with knowledge of the odor sequence, suggesting this signal is associated with learning, and was differentially recruited across trial types, offering strong evidence for its behavioral relevance. Second, we found that 20-40 Hz power was higher in proximal than distal CA1, which is the opposite pattern to that observed in an odor-place association task (Igarashi et al., 2014), suggesting that oscillatory power gradients along the proximodistal axis may reflect task-specific demands. In light of prior evidence that proximal CA1 is strongly associated with the medial entorhinal cortex (MEC; van Strien, Cappaert, and Witter, 2009; Witter et al., 2017), and that MEC inactivations impair temporal coding in CA1 (Robinson et al., 2017), this finding suggests that functional coupling between proximal CA1 and MEC may play a key role in remembering the temporal context of nonspatial events.

**Figure 1.**
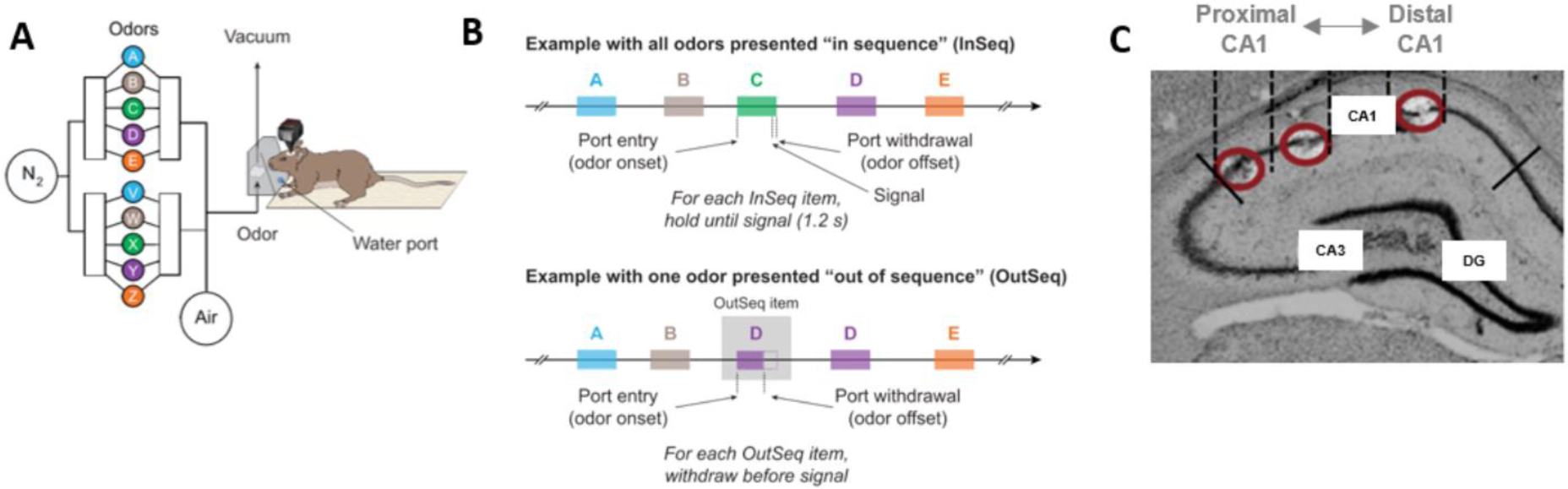
Odor sequence task and electrode locations. **A.** Using an automated odor delivery system, rats were presented with series of five odors delivered in the same odor port (located at one end of a linear track). **B.** In each session, the same sequence was presented multiple times, with approximately half the presentations including all items “in sequence” (InSeq; ABCDE) and the other half including one item “out of sequence” (OutSeq; e.g., ABDDE). Each odor presentation was initiated by a nosepoke and rats were required to correctly identify the odor as either InSeq (by holding their nosepoke response until the signal at 1.2 s) or OutSeq (by withdrawing their nosepoke before the signal; <1.2 s) to receive a water reward. After completion of each sequence (correctly or incorrectly), animals were required to run to the other end of the linear track and return to the odor port before the next sequence could be presented. **C**. Sample histology image showing the range of tetrode tip locations, which spanned much of the proximodistal axis of CA1 (3 tip locations shown; red circles). For each animal, a set of four tetrodes equally distributed across the proximodistal axis (with comparable locations across animals) was used for local field potential activity analysis.

## Results

### Oscillatory states differ between odor sequence processing and running periods and vary across the proximodistal axis of CA1

We began by investigating whether there are distinct oscillatory states in CA1 that are unique to the odor sequence processing component of the task, and whether the observed oscillations varied across the proximodistal axis of CA1. To do so, group (n=5) peri-event spectrograms were generated from four electrode locations along the proximodistal axis and aligned to odor processing (Fig 2A) and running (Fig 2B) epochs. Data were taken from a session in which animals performed at a high level (well-trained session). We observed that power in the 20-40 Hz and 4-8 Hz range observed during odor presentations showed significantly distinct patterns across the proximodistal axis (Fig 2A,C,D; Electrode x Band interaction: F_3,12_ = 8.995, *p* = 0.0021). Power of 20-40 Hz oscillations increased toward proximal CA1 (One-way ANOVA: F_3,12_ = 4.4184, *p* = 0.0284; Linear trend across electrodes: F_1,12_ = 10.56, *p* = 0.0070; Individual subjects ANOVAs: significant in 4 out of 5 subjects, see Table S1A). In contrast, 4-8 Hz power was numerically higher in distal CA1 although the one-way ANOVA and linear trend analysis did not reach significance, possibly due to an outlier (one-way ANOVA: F_3,12_ = 1.0237, *p* = 0.4165; Linear trend across electrodes: F_1,12_ = 2.434, *p* = 0.1447; Individual subjects ANOVAs: significant in 4 out of 5 subjects, with the fifth subject showing opposite pattern; see Table S1B).

**Figure 2.**
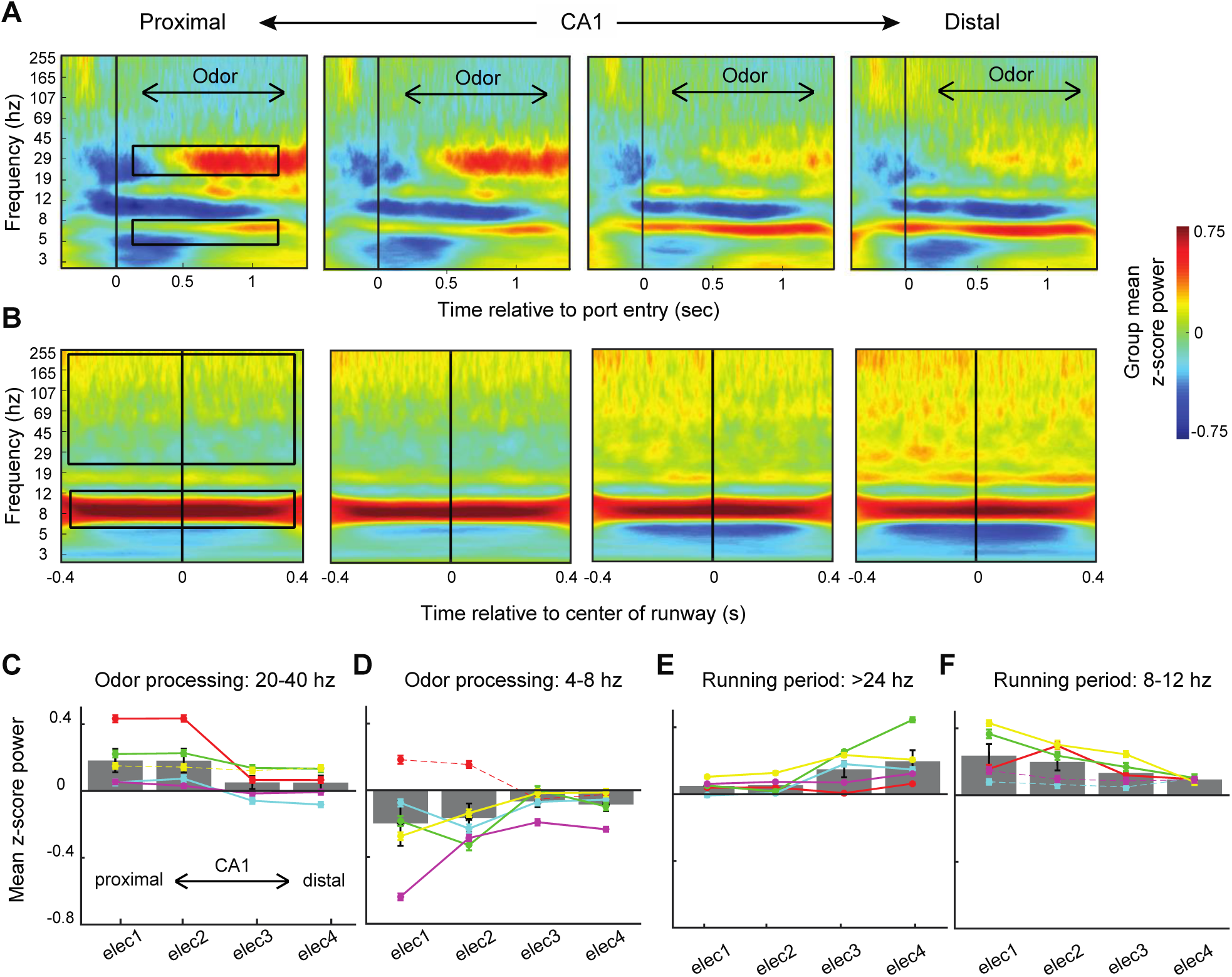
Odor sequence processing and running on a track are associated with distinct oscillatory states in CA1, which vary across the proximodistal axis. **A.** Group peri-event spectrograms (n=5) during odor sampling period (correct in sequence trials only) in four electrode locations along the CA1 proximodistal axis (0ms = port entry). **B.** Group peri-event spectrograms from the same electrodes during the running period (0ms = center of the runway).**C.** Mean z-score power for 20-40 Hz oscillations during 110-1200ms period of odor presentation (time period indicated by upper black box in panel A; defined *a priori*). **D.** Mean z-score power for lower frequency (odor-associated) theta oscillations (4-8 Hz) during 110-1200ms period of odor presentations (indicated by lower black box in panel A). **E.** Mean z-score power for higher frequency oscillations (>24 Hz to avoid theta’s first harmonic) during running period (indicated by upper black box in panel B). **F.** Mean z-score power for higher frequency (running-associated) theta oscillations (8-12 Hz) during running period (indicated by lower black box in panel B). Both odor sampling and running intervals were extracted from a session in which animals performed at a high level (well-trained session). For each electrode site, spectrograms were generated using analytic Morlet wavelets and spectral power was z-scored relative to the power at the same site during a 30-minute period of the recording session (negative values represent mean power values below baseline). Grey bars indicate group means (error bars represent SEM across subjects). Colored circles indicate means for individual subjects (error bars represent SEM across trials), with solid and dashed lines representing significant (*p* < 0.05) and non-significant individual subjects ANOVAs, respectively. See Tables S1-S2 for statistical results.

The same electrodes exhibited a different pattern of oscillations during running periods (which occurred between sequence presentations; Fig 2B,E,F). More specifically, the 20-40 Hz and 4-8 Hz bands observed during odor sampling were weak during running. Instead, we observed strong oscillations in the 8-12 Hz theta range during running (consistent with numerous reports), which increased toward proximal CA1 (one-way ANOVA: F_3,12_ = 3.0296, *p* = 0.0711; Linear trend across electrodes: F_1,12_ = 8.944, p = 0.0113; Individual subjects ANOVAs: significant in 3 out of 5 subjects, see Table S2B). The running period was also characterized by increased power in higher frequencies (> 24 Hz to avoid theta’s first harmonic), which increased toward distal CA1 (one-way ANOVA: F_3,12_ = 3.7614, *p* = 0.0410; Linear trend across electrodes: F_1,12_ = 10.21, p = 0.0077; Individual subjects ANOVAs: significant in 5 out of 5 subjects, see Table S2A). Notably, the proximodistal pattern was significantly different between the two bands (Electrode X band interaction: F_3,12_ = 5.343, *p* = 0.0144).

### 20-40 Hz power increases with knowledge of the sequence

We then examined whether power in the 20-40 Hz range was linked with sequence memory performance, and whether this association varied along the proximodistal axis. To do so, we extended the previous analyses, which were applied to a well-trained session, to two consecutive sessions in which animals learned a novel odor sequence (correctly identified InSeq trials only). This allowed us to compare power across three sessions characterized by low (first session on novel sequence), moderate (second session on novel sequence) and high levels of performance (well-trained session; Fig 3). Unlike the previous analyses, here we evaluated 20-40 Hz power during the 250ms time window preceding the port withdrawal response. This was done to capture the period of high power observed toward the end of odor presentations and to provide the same alignment as the trial-type analyses described in the next section. This approach yielded higher power values than the previous analysis (compare Fig 3B with Fig 2C) but confirmed the pattern in which 20-40 Hz power increases toward proximal CA1 (one-way ANOVA: F_3,12_ = 8.0838, *p* = 0.0032; Individual subjects ANOVAs: significant in 4 out of 5 subjects, see Table S3A).

**Figure 3.**
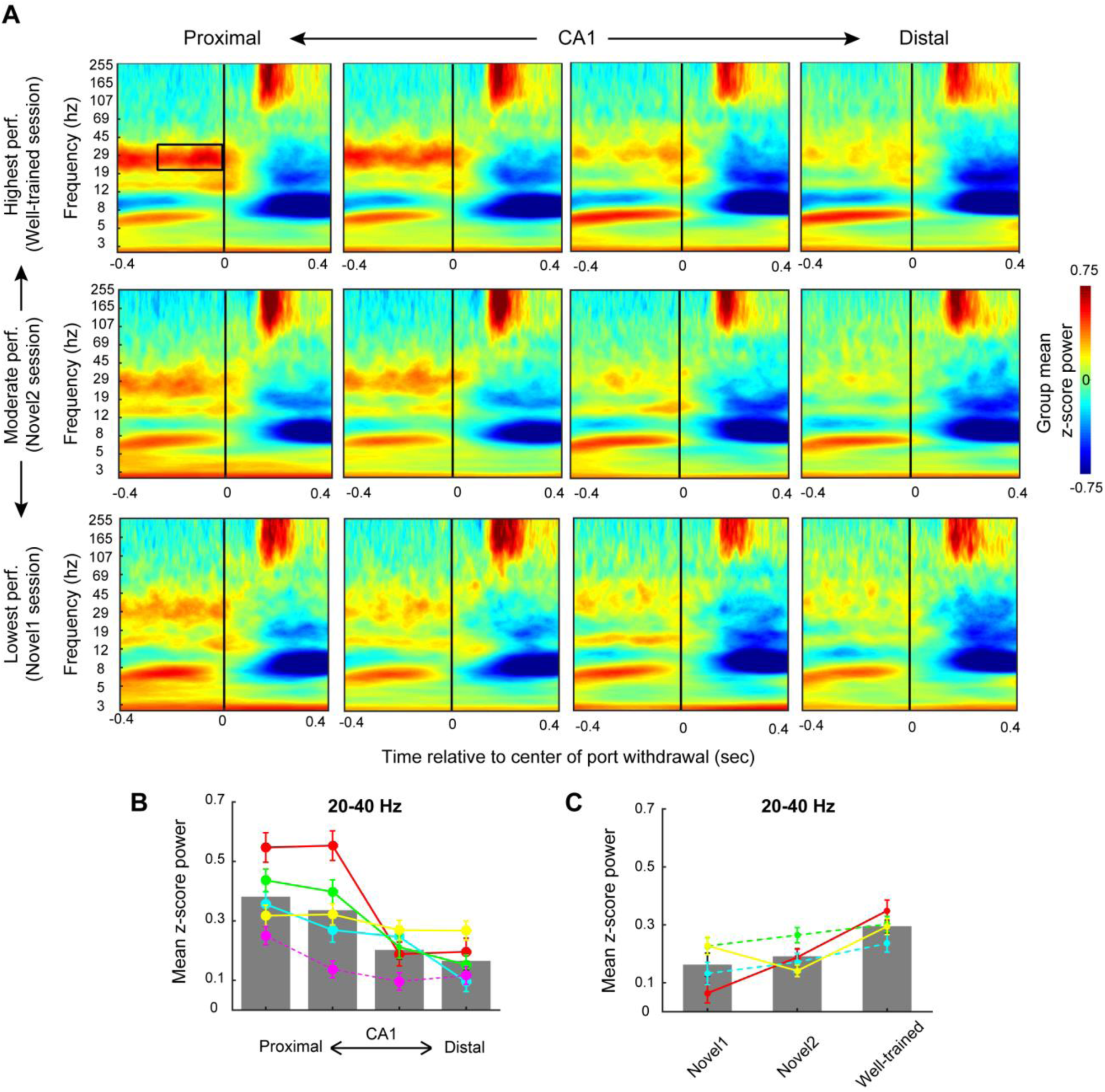
CA1 20-40 Hz power increases with knowledge of the sequence. **A.** Group peri-event spectrograms across three sessions in which performance levels were low (first session on novel sequence; *Bottom row*), moderate (second session on novel sequence; *Middle row*) and high (well-trained session; *Top*). All plots are aligned to port withdrawal (0ms = port withdrawal) and only include correctly identified InSeq trials. **B-C**. Mean z-score power in the 250 ms period prior to port withdrawal (indicated by black box in top left panel) across the four proximodistal electrode sites (**B**) and three sessions with increasing performance levels (**C**). Grey bars indicate group means (error bars represent SEM across subjects). Colored circles indicate means for individual subjects (error bars represent SEM across trials), with solid and dashed lines representing significant (*p* < 0.05) and non-significant individual subjects ANOVAs, respectively. See Table S3 for statistical results.

We found that 20-40 Hz power increased with performance level (Fig 3C; one-way ANOVA: F_2,6_ = 5.51, *p* = 0.0438; Individual subjects ANOVAs: significant in 2 out of 4 subjects; see Table S3B), which complements our previous study showing learning-related differences in waveform amplitude between InSeq and OutSeq trials (Allen et al., 2016). In addition, we found that this performance effect did not significantly vary across the proximodistal axis, but instead scaled with the local amplitude of the 20-40 Hz oscillation, suggesting this signal is present throughout dorsal CA1 (data not shown).

### 20-40 Hz power varies with response type and accuracy

To shed light on the type of processing reflected by 20-40 Hz oscillations, we took advantage of the four different trial types included in our paradigm: InSeq trials that were correctly or incorrectly identified (InSeq+, InSeq-), and OutSeq trials correctly or incorrectly identified (OutSeq+, OutSeq-). More specifically, we quantified power in the 20-40 Hz range (250ms period before port withdrawal; averaged across the four electrodes) for each trial type separately, as well as collapsing across trial types to test contrasts of particular interest (Fig 4). To match previous plots, the data are first presented using a single value per animal to generate the group mean (Fig 4B), despite the increased variability induced by trial count discrepancies across trial types (see Fig S1B and Table S4 for trial counts). To control for this, the data are also presented using an approach in which we pooled trials from all animals and used a sampling procedure to match trial count across trial types (Fig 4C).

**Figure 4.**
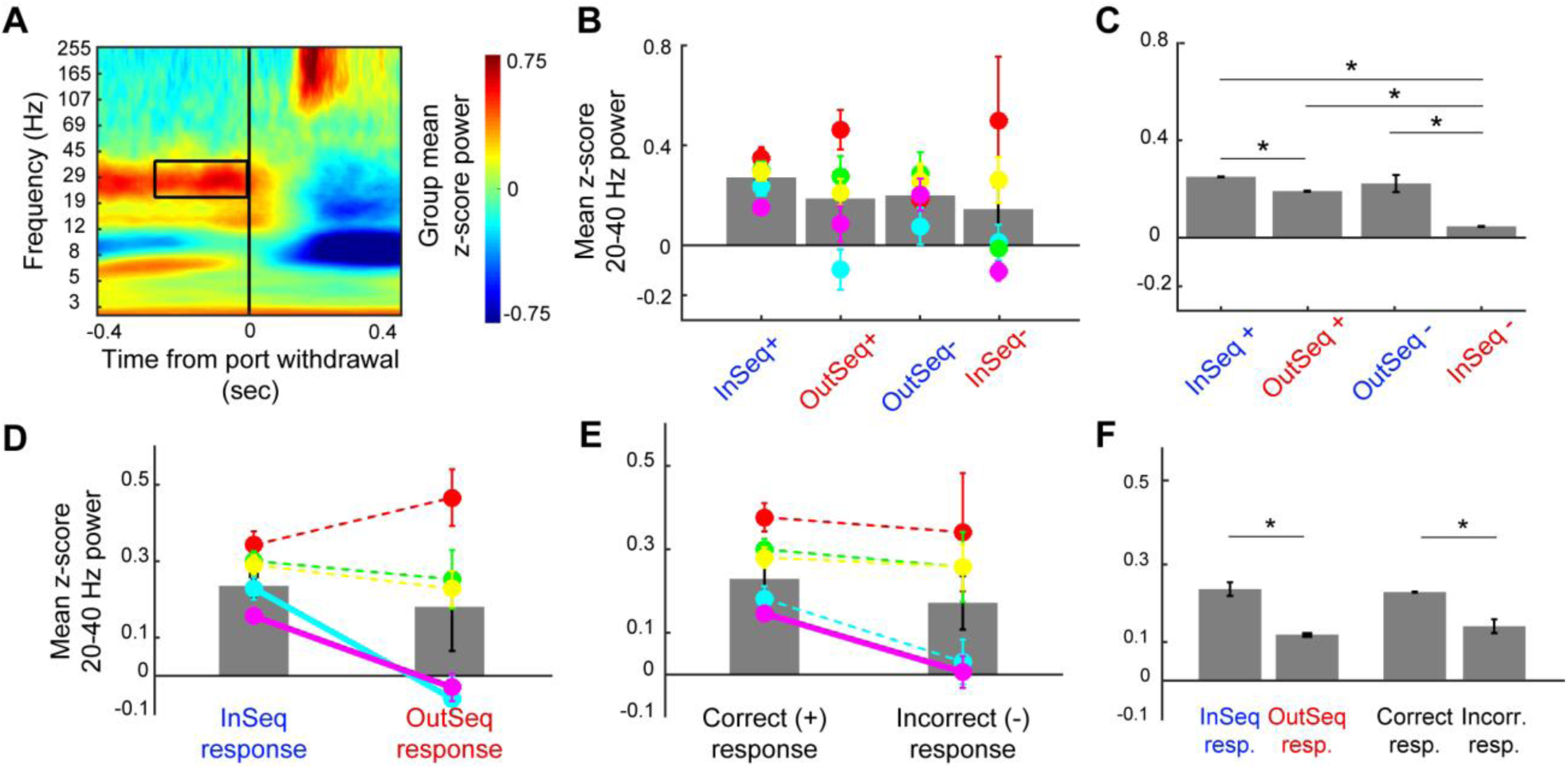
CA1 20-40 Hz power varies across trial types. **A.** Group peri-event spectrogram relative to port withdrawal during the well-trained session (most proximal CA1 site). **B.** Mean z-score 20-40 Hz power (250ms period preceding port withdrawal) across four trial types: InSeq trials that were correctly or incorrectly identified (InSeq+, InSeq-), and OutSeq trials correctly or incorrectly identified (OutSeq+, OutSeq-). To match previous plots, data are presented using one value per subject, despite known differences in trial count across trial types. Grey bars indicate group means (error bars: SEM across subjects) and colored circles indicate means for individual subjects (error bars: SEM across trials). **C.** Same as in B, with the exception that averaging is performed across trials pooled from all animals using a sampling procedure to match trial count across trial types (error bars indicate SEM across pooled trials). **D.** Contrast between InSeq responses (InSeq+ and OutSeq-trials) and OutSeq response (InSeq- and OutSeq+ trials) using one value per animal. **E.** Contrast between correct (InSeq+ and OutSeq+) and incorrect (InSeq- and OutSeq-) trials using one value per animal. **F.** Same contrasts as in D and E, using sampling procedure to match trial count across trial types. * significant permutation test with FDR correction for multiple comparisons (all significant corrected *p* values are <.002). See Table S4 for trial counts.

First, to test whether 20-40 Hz oscillations reflect a match/mismatch signal between the stimulus *presented* and the stimulus *predicted* by the animal (based on its knowledge of the sequence), we directly compared power between InSeq+ and OutSeq+ trials. InSeq+ trials represent a matched prediction (e.g., in the second sequence position, the animal predicted B and was presented with B), whereas OutSeq+ represents a mismatched prediction (e.g., in the second position, the animal predicted B but was presented with C). We confirm that InSeq+ trials showed significantly higher 20-40 Hz power than OutSeq+ trials (*p* = 0.002 using permutation testing with FDR correction for multiple comparisons; see methods), consistent with previous findings using waveform amplitude (Allen et al., 2016). However, it is important to note that this *a priori* contrast is confounded by the *posthoc* observation that 20-40 Hz power gradually increases during odor presentations, in that power may be higher on InSeq+ trials because their corresponding time window occurred later in the trial than OutSeq+ trials. Therefore, this effect will require confirmation using a different paradigm.

Second, to test whether 20-40 Hz oscillations reflected the type of response produced, we compared power between InSeq responses (InSeq+ and OutSeq-trials) and OutSeq responses (InSeq- and OutSeq+ trials). Using the pooled and sampled trial distributions (to balance trial count across trial types) we found that power in the 20-40 Hz range was significantly higher on InSeq responses than OutSeq responses (*p* = 0.002, permutation testing; Fig 4F). As expected, the effect was in the same direction, but considerably more variable, when only considering a single value per animal (Fig 4D). Third, to test whether these oscillations are associated with accurately performing the cognitive operations required on each trial, we compared power between correct (InSeq+ and OutSeq+) and incorrect (InSeq- and OutSeq-) trials. Power was significantly higher on correct trials (*p* = 0.002, permutation testing; Fig 4F). As above, the effect was more variable but in the same direction when considering only one value per subject (Fig 4E). It is important to note that the effects on the latter two contrasts (InSeq vs OutSeq responses; Correct vs Incorrect trials) were primarily driven by the fact that InSeq-trials showed the lowest level of 20-40 Hz power of all trial types. This suggests that information about both response type and accuracy are reflected in these oscillations, though the relative degree of their contributions remains unclear.

### 20-40 Hz power increases are not simply due to increased response duration

As shown in Figure 2, power in the 20-40 Hz range increases toward the end of the odor presentations. This raises the possibility that, in addition to the cognitive processes described in the previous section, power increases may be linked to the duration of the nosepoke response. This could be due to the increased demand of sustaining posture in the port for an extended time or it could reflect a slow-rising signal that reaches plateau on trials in which long nosepoke responses were performed. We tested this possibility using two approaches. First, we examined whether there was a linear relationship between nosepoke duration and 20-40 Hz power across trials for each of the 20 electrodes included in the previous analyses. We focused this analysis specifically on InSeq+ trials to match cognitive demand and maximize statistical power (InSeq+ trials have the highest count). We found that correlations were near zero for the majority of electrodes (Fig S1A), with only one electrode (out of 20) showing a significant correlation (r = 0.2722, *p* = 0.0009). Second, we examined whether the pattern of response durations matched the pattern of 20-40 Hz power across trial types. We found that this was not the case (compare Fig S1C with Fig 4C), indicating that the observed power dynamics across trial type could not have arose simply due to differences in response time. For instance, 20-40 Hz power was not significantly different between OutSeq+ and OutSeq-, but response durations were considerably shorter on OutSeq+ trials (mean duration: ∼750ms) than OutSeq-trials (mean duration: ∼1,500ms). In addition, InSeq-trials displayed the lowest 20-40Hz power despite most responses occurring near the decision threshold, considerably later than OutSeq+ responses (InSeq-mean duration: ∼975ms; compare with ∼750ms for OutSeq+; Fig S1B,C). Collectively, these analyses demonstrate that 20-40 Hz power dynamics are not simply driven by response duration.

### Gamma power is not differentially associated with trial type-specific information

Finally, given the prior work on slow and fast gamma in spatial navigation tasks, we assessed whether these ranges were associated with specific epochs of the task, supporting memory for the temporal order of events. As previously demonstrated in spatial navigation tasks (Colgin et al., 2009), we observed power increases in the slow gamma (25-55 Hz) and high gamma (60-100 Hz) bands during running epochs (most noticeably in distal CA1; Fig 2B). However, neither slow (25-55 Hz) nor fast (60-100 Hz) gamma oscillations showed distinctive trial type-specific information in the putative retrieval (250 ms prior to port entry) and encoding (110-300 ms following port entry) windows, respectively (Fig S2). However, we note that the slow gamma range (25-55 Hz) overlaps with the observed 20-40 Hz oscillations prior to port withdrawal, which may be a putative retrieval period. Overall, these findings suggest that the pattern of distinct slow gamma and fast gamma oscillatory states observed in CA1 during spatial navigation may be not be readily visible in the predicted epochs of a nonspatial sequence processing task. However, it is possible that our experimental and analytical approach were not optimal to directly test these effects.

## Discussion

In this paper, we examined oscillatory power in hippocampal region CA1 as rats performed a complex sequence memory task to identify the oscillatory dynamics associated with nonspatial information processing. The data presented here expand upon our previous report of 20-40 Hz oscillations during the odor sequence processing periods of the task by evaluating the behavioral relevance and spatial distribution of such oscillations across the CA1 proximodistal axis, as well as providing a direct comparison with another behavioral state (running). First, we demonstrate that running and odor sequence processing epochs are characterized by different spectral content. Running is associated with increased power in 8-12 Hz and >24 Hz ranges, whereas odor sequence processing is associated with increased power in 4-8 Hz and 20-40 Hz ranges. Second, we show that in both behavioral states, there are significantly distinct gradients with respect to power of the recruited oscillations along the CA1 proximodistal axis. During odor processing epochs, 20-40Hz power is higher in proximal CA1, whereas 4-8 Hz power is numerically (but not significantly) higher in distal CA1. During running periods, 8-12 Hz power is higher in proximal CA1 whereas >24 Hz power is higher in distal CA1. Third, we found that the 20-40 Hz oscillation is linked with sequence memory performance. Power in this range increases with session performance and varies across trial types. More specifically, 20-40 Hz power is higher for trials with an “in sequence” response (a presumed match between the presented odor item and the predicted one) and during correct compared to incorrect trials. Lastly, we do not find evidence that slow and fast gamma oscillations previously observed during spatial exploration tasks are associated with specific trial types during the putative encoding and retrieval epochs of the task, although we did not test other epochs for this effect. We suggest that more work needs to be done to fully ascertain the role of slow and fast gamma oscillations in nonspatial tasks. Altogether, these findings suggest that processing the temporal context of nonspatial events primarily recruits oscillations in the 20-40 Hz range in the proximal segment of CA1.

It is important to note that the nature of this complex experimental paradigm led to two potential limitations to consider when interpreting the findings. First, the use of a nonspatial response (hold/withdraw) prevented us from directly equating response duration across trial type. Although the degree to which differences in response duration influence our findings cannot be fully determined, we showed that response duration alone does not explain the differential recruitment of 20-40 Hz power across trial types. Second, to ensure adequate performance, the task requires that the number of OutSeq trials be kept relatively low (otherwise it is unclear which sequence is being tested). This results in an uneven number of observations across trial types which could disproportionally influence a subset of our analyses. However, we controlled for this possibility by conducting pooled analyses, which matched trial count across conditions using a permutation sampling procedure, to ensure sufficient statistical power. Overall, we believe these control analyses significantly mitigated the potential influence of these confounding factors on the interpretation of our results.

Prior studies have shown a similar recruitment of 20-40 Hz oscillations in odor-based tasks. Oscillations in a similar range (15-35 Hz) were recorded in the hippocampus and olfactory bulb of rats performing a go/no-go odor discrimination task (Martin et al., 2007). Interestingly, that study showed a learning-related increase in oscillatory power in the olfactory bulb as well as in the coherence between signals from bulb and hippocampus. However, oscillatory power in the hippocampus did not change as a function of learning. Similarly, 20-40 Hz oscillations were observed in distal CA1 (dCA1) and lateral entorhinal cortex (LEC) of rats engaged in an odor-place association task (Igarashi et al., 2014). In that study, learning-related increases in dCA1-LEC coherence were observed, but oscillatory power in either region did not significantly change with learning. This suggest that although these oscillations are observed in a variety of odor-based tasks, increases in odor familiarity or task performance do not necessarily result in a corresponding increase in power. Instead, 20-40 Hz power increases may be linked to specific task demands, such as the processing of the temporal context of events.

Evidence for functional heterogeneity along the CA1 proximodistal axis has been previously reported (Henriksen et al., 2010; Hartzell et al., 2013; Nakazawa et al., 2016; Ng et al., 2018). Such heterogeneity is not too surprising given the known gradients of connectivity from the lateral and medial entorhinal cortices (LEC and MEC; van Strien, Cappaert, and Witter, 2009; Witter et al., 2017), specifically the fact that distal CA1 is more strongly associated with LEC, both anatomically and functionally, whereas proximal CA1 is more strongly associated with MEC. Consequently, the observation of different proximodistal gradients of oscillatory power across studies may result from differences in task demands, which could promote differential engagement of these entorhinal-CA1 circuits (although other factors, including electrode placements and analytical approaches, may also contribute to this difference). For instance, Igarashi et al (2014) described that 20-40 Hz power was higher in *distal* CA1 during performance of an odor-place association task (whereas we showed higher power in *proximal* CA1). Since performance in their task depends on the correct identification of the specific perceptual features that distinguish one odor from another, the power increase in distal CA1 may reflect a stronger engagement of the LEC-dCA1 component of the circuit (as LEC receives strong olfactory input, including direct projections from the olfactory bulb; Haberly and Price, 1978; Agster and Burwell, 2009). In contrast, in our experiment, identification of the presented odor is alone insufficient for correct performance --- the animal must further identify whether or not the odor is being presented in the appropriate temporal context. The power increase in proximal CA1 we observed may reflect the fact that this additional temporal requirement preferentially engaged the MEC-pCA1 component of the circuit. Interestingly, although MEC is typically associated with spatial navigation functions, this interpretation is supported by a recent study by Robinson and colleagues (2017), in which they demonstrated that optogenetic inactivation of MEC disrupted temporal coding in CA1, while sparing spatial and object coding. Together with the proximodistal pattern we observed, these results suggest that perhaps the MEC→proximal CA1 microcircuit may be important for the processing of nonspatial temporal information.

While the evidence reported here suggests a role for 20-40 Hz oscillations in proximal CA1 in processing nonspatial information, the origin of this rhythm remains unclear. CA1 receives input from a number of other sources including EC, CA2, CA3, and the medial septum (van Strien, Cappaert, and Witter, 2009). These upstream structures may contribute to the generation of this oscillation in CA1, though it may also be locally generated within CA1. It is also worth noting that the 20-40 Hz frequency range prominent here overlaps with the previously reported slow gamma band (25-55 Hz), which has been implicated in memory retrieval, shown to be coherent with CA3, and is thought to be involved in the routing of information from CA3 to CA1 (Colgin et al., 2009). It is therefore possible our findings on 20-40 Hz power include contributions from slow gamma, or that the two oscillations reflect overlapping mechanisms. Further studies with multisite recordings will be required to assess these possibilities.

As with other neural recording data, it is difficult to determine what the observed 20-40 Hz power dynamics reflect in terms of information processing and how they are linked with behavior. Oscillations in this range (typically referred to as beta) occur widely across the cortex and have been associated with different functions across brain regions (Engel & Fries, 2010; Schmidt et al., 2019). Of particular relevance here, beta has been associated with olfactory processing (Martin & Ravel, 2014), temporal estimation (Wiener, Parikh, Krakow & Coslett, 2018), working memory (Miller, Lundqvist & Bastos, 2018), and postural maintenance (Kilavik et al., 2013). However, these accounts do not fully capture the complexity and specificity of the power dynamics we observed in the present study. Our findings that 20-40 Hz power increases with learning, is higher on InSeq responses and on correct trials, suggest this signal is associated with trial-specific computations critical to solve the task. The observation that power gradually increases during odor presentations and abruptly decreases after the port withdrawal response is consistent with this is well. One possibility is that 20-40 Hz power reflects a degree of match between the stimulus presented and the stimulus predicted by the animal (based on its knowledge of the sequence). This possibility is well aligned with the hypothesized role of CA1 acting as a comparator between internal representations retrieved from the CA3 and external cues transmitted via the entorhinal cortex (e.g., Hasselmo and Wyble, 1997; Lisman and Grace, 2005) and is consistent with the strong InSeq/OutSeq differentiation observed in spiking activity at the single-cell (Allen et al., 2016) and ensemble (Shahbaba et al., 2019) level. The learning-related power increase we observed is also consistent with this view, as stimulus predictions should improve with learning (resulting in stronger matches on InSeq+ trials). The observation that power is higher on InSeq+ than OutSeq+ trials would also be consistent with this view, but this effect would need to be confirmed using a paradigm in which response duration can be matched across trial types. Finally, it is important to consider the possibility that the highest 20-40 Hz power values observed near the end of stimulus presentations, in our paradigm and that of others, may reflect a post-decision state (OutSeq+ responses were, on average, made ∼750ms after port entry). Thus, power increases observed earlier in the trial may be more strongly linked to information processing steps leading to behavioral decisions.

In conclusion, our demonstration of learning-related and trial-specific increases in 20-40 Hz power links this oscillation with task-critical information processing. Future work will be needed to identify the generator of this rhythm and the specific cognitive processes or computations reflected by this signal.

## Methods

Our group previously published using the same dataset, and a detailed description of the methods can be found in Allen and colleagues (2016). The methods are summarized below.

### Subjects

Five male Long-Evans rats were used in this study. Animals were water restricted for optimum task engagement but were provided full access to water on weekends. Proper hydration levels were monitored throughout the experiment. All procedures were conducted in accordance with the guidelines from care and use of laboratory animals published by the National Institutes of Health. All animals were handled according to approved Institutional Animal Care and Use Committee (IACUC) protocols. Sample sizes were determined using standards in behavioral electrophysiology experiments. Data was recorded from 5 animals (each animal represents several months of work), with each animal providing data from 20 electrodes over a minimum of 103 trials. In total, the dataset included 100 electrodes and 785 trials.

### Replicates

Although it takes several months to train, implant, and record from each animal, the “experiment” focused on three daily sessions per animal (matched across animals). Data from the same animal was <u>not</u> collapsed across sessions. In our design, we view animals as biological replicates and, within each animal, the number of trials as technical replicates. Electrodes can be viewed as biological replicates (e.g., when comparing effects across electrodes within each animal) or technical replicates (e.g., when collapsing across electrodes to confirm a general pattern was present across electrodes). The supplementary tables included provide detailed information on the number of trials included in each statistical comparison. The number of animals is included in the main text (p14) and in the supplementary tables.

### Outliers and Inclusion/Exclusion of Data

No statistical outliers were removed. Standard pre-processing approaches were used to exclude data contaminated by electrical noise or artifacts. As stated in the manuscript (page 16), 60Hz electrical noise was removed using a notch filter. Trials with artifacts associated with bumping or touching the headstage (voltage values > 5 SD above the mean) were automatically excluded. Note that this exclusion was performed before (and blind to) analysis of the results.

### Equipment

The apparatus used for this task consisted of a linear track with water ports on either end for water reward delivery. One end of the maze contained an odor port (above the water port) connected to an automated odor delivery system. Photobeam sensors detected when the animal’s nose entered and withdrew from the odor port, which respectively triggered and terminated odor delivery. Separate tubing lines were used for each odor item, however, all converged at a single channel at the bottom of the odor port. The odor port was kept clear of previous odor traces using a negative pressure vacuum located at the top of the port. A 96-channel Multichannel Acquisition Processor (MAP; Plexon) was used to interface the hardware (Plexon timing boards and National Instruments input/output devices) in real time and record the behavioral and electrophysiological data as well as control the hardware.

### Odor sequence task

In this hippocampus-dependent task, rats were presented with series of five odors delivered in the same odor port (Fig 1). In each session, the same sequence was presented multiple times, with approximately half the presentations including all items “in sequence” (InSeq; ABCDE) and the other half including one item “out of sequence” (OutSeq; e.g., AB<u>D</u>DE). Each odor presentation was initiated by a nosepoke and rats were required to correctly identify the odor as either InSeq (by holding their nosepoke response until the signal at 1.2 s) or OutSeq (by withdrawing their nosepoke before the signal; <1.2 s) to receive a water reward. Animals were trained preoperatively on sequence ABCDE (lemon, rum, anise, vanilla, and banana) until they reached asymptotic performance (>80% correct on both InSeq and OutSeq trials; ∼6 weeks). Following surgical recovery, electrophysiological data was collected as animals performed the same sequence (ABCDE), followed by two consecutive sessions using a novel sequence (VWXYZ; almond, cinnamon, coconut, peppermint, and strawberry).

### Surgery

Rats received a preoperative injection of the analgesic buprenorphine (0.02 mg/kg, s.c.) ∼10 min before induction of anesthesia. General anesthesia was induced using isoflurane (induction: 4%; maintenance: 1–2%) mixed with oxygen (800 ml/min). After being placed in the stereotaxic apparatus, rats were administered glycopyrrolate (0.5 mg/ kg, s.c.) to help prevent respiratory difficulties. A protective ophthalmic ointment was then applied to their eyes and their scalp was locally anesthetized with marcaine (7.5 mg/ml, 0.5 ml, s.c.). Body temperature was monitored and maintained throughout surgery and a Ringer’s solution with 5% dextrose was periodically administered to maintain hydration (total volume of 5 ml, s.c.). The skull was exposed following a midline incision and adjustments were made to ensure the skull was level. Six support screws (four titanium, two stainless steel) and a ground screw (stainless steel; positioned over the cerebellum) were anchored to the skull. A piece of skull ∼3 mm in diameter (centered on coordinates: -4.0 mm anteroposterior, 3.5 mm mediolateral) was removed over the left hippocampus. Quickly after the dura was carefully removed, the base of the microdrive was lowered onto the exposed cortex, the cavity was filled with Kwik-Sil (World Precision Instruments), the ground wire was connected, and the microdrive was secured to the support skull screws with dental cement. Each tetrode was then advanced ∼900 μm into the brain. Finally, the incision was sutured and dressed with Neosporin and rats were returned to a clean cage, where they were monitored until they awoke from anesthesia. One day following surgery, rats were given an analgesic (flunixin, 2.5 mg/kg, s.c.) and Neosporin was reapplied to the incision site.

### Electrophysiological recordings

Both spiking and local field potential activity were recorded from the CA1 pyramidal layer of the dorsal hippocampus as rats performed the task (see Allen et al., 2016), but the present study focuses exclusively on a detailed analysis of the LFP activity. Each chronically implanted microdrive contained 20 independently drivable tetrodes, with each tetrode consisting of four twisted nichrome wires (13 μm in diameter; California Fine Wire) gold-plated to achieve a final tip impedance of ∼250 kΩ (measured at 1 kHz). Following the surgical recovery period, tetrodes were slowly advanced over a period of ∼3 weeks while monitoring established electrophysiological signatures of the CA1 pyramidal cell layer (e.g., sharp waves, ripples, and theta amplitude). Voltage signals from electrode tips were referenced to a ground screw positioned over the cerebellum. LFP activity was filtered (1.5 - 400 Hz), amplified (1000X), digitized (1 kHz), and recorded to disk with the data acquisition system (MAP, Plexon). Neural activity data was first recorded on the odor sequence learned before surgery (ABCDE; “Well-trained” session), followed by two consecutive sessions on the same novel sequence (VWXYZ; Novel1 and Novel2 sessions). At the end of the experiment, recording sites were confirmed by passing current through the electrodes before perfusion (0.9% PBS followed by 4% para-formaldehyde) to produce small marking lesions, which were subsequently localized on Nissl-stained tissue slices.

### Preprocessing and spectral analysis

The raw data was pre-processed using a Butterworth notch filter to remove 60 Hz line noise. Artifact rejection was defined by time indices with time domain voltage values greater than 5 standard deviations above the mean signal of the entire recording in the same channel. Artifact time points were included for the wavelet processing in order to maintain the temporal structure of the data, but their associated power values were removed before computing the mean and standard deviation of baseline used for normalization (see below). Any trial containing an artifact was excluded from analyses. For spectral analysis, we utilized the Wavelet toolbox in MATLAB (Mathworks) to generate analytic Morlet wavelets for frequencies between 3 to 250 Hz. These wavelets were tested and verified on a simulated data with known spectral properties. Next, we extracted behavior-locked instantaneous power at the specified frequency ranges. In all analyses, the first trial of each sequence was excluded as it was always preceded by running, whereas the animal was stationary prior to all other trial positions.

### Normalization

Instantaneous power is reported as a z-score value relative to the mean and standard deviation of power for a given frequency calculated from a 30-minute subset of the recording from the same electrode. For comparison, we also used two additional normalization approaches. One approach calculated z-scores relative to the other time points within the same trial (0 to 1.5 s for trials aligned to port entry; -1s to 0s for trials aligned to port withdrawal). In the other approach, power value for a given time point and frequency within a trial were divided by the sum of the power across all trial time points in the same frequency, which captured percentage increase in power at a given frequency. As all three methods yielded comparable results the reported results relied on the z-normalization to the 30-minute recording subset. As this 30-minute period included a variety of behavioral and cognitive states, including odor sampling, running, grooming, and reward consumption, it offers a better characterization of the variance of oscillatory dynamics associated with the animals’ experiences.

### Selection of electrodes along the proximodistal axis

In order to sample four representative electrodes along the proximodistal axis of CA1, we chose the first and the last electrodes (most proximal and most distal, respectively) and two electrodes in between which were equidistant. We confirmed the relative spatial distribution of these electrodes, as well as their localization within the pyramidal layer of CA1 based on standard spectral properties during baseline, odor sampling, and running periods. For each of the four electrodes selected per animal, LFP activity patterns were confirmed in adjacent electrodes (from the remaining subset of 16 electrodes).

### Sampling procedure for comparisons across trial types

Analyses comparing across trial types used a sampling procedure to account for disparities in the number of trials (see Fig 4C; Table S4). Trials were first pooled across all animals and the condition with the minimum trial count was identified (e.g. OutSeq-, n = 47). Then, in the remainder conditions (InSeq+, InSeq-, and OutSeq+), 47 trials were randomly chosen 1000 times. This generated three distributions, one for each condition, of randomly sampled 47 trials. This sampling procedure enabled for sufficient statistical power to examine group effects in conjunction with statistical examination on an individual animal basis.

### ANOVA and permutation testing

Group analyses were performed using one-way and two-way ANOVAs with repeated measures, followed up with linear trend analyses (Prism 8.0). Individual subjects’ one-way ANOVAs were performed in MATLAB (anova1 function), followed up by pair-wise permutation testing with FDR correction (see tables associated with each figure). Permutation testing was also used for group analyses involving pair-wise comparisons across trial types. Permutation testing was performed by shuffling trial labels 1,000 times, with the *p* value representing the probability of obtaining a mean difference as high (or higher) than the one observed through random shuffling.

### Data availability

Data are available on Dryad (doi: 10.7280/D11960) and code/scripts used to generate all paper figures and reported statistics are available on Github (https://github.com/FortinLab/Gattas_et_al_2020).

## Acknowledgments

This research was supported in part by the National Science Foundation (awards IOS-1150292 and BCS 1439267 to N.J.F.), the National Institutes of Health (awards R01 MH115697 and R01 DC017687 to N.J.F.; R01 MH102392 and R01 AG053555 to M.A.Y.; T32 NS45540 support for S.G.; and T32 DC010775 support for G.A.E.), and the Whitehall Foundation (award 2010-05-84 to N.J.F.). We would also like to thank Aaron Gudmundson and members of the Fortin Lab for useful discussions on the present work.

## Supplementary Materials

### Supplementary Figures

**Figure S1.**
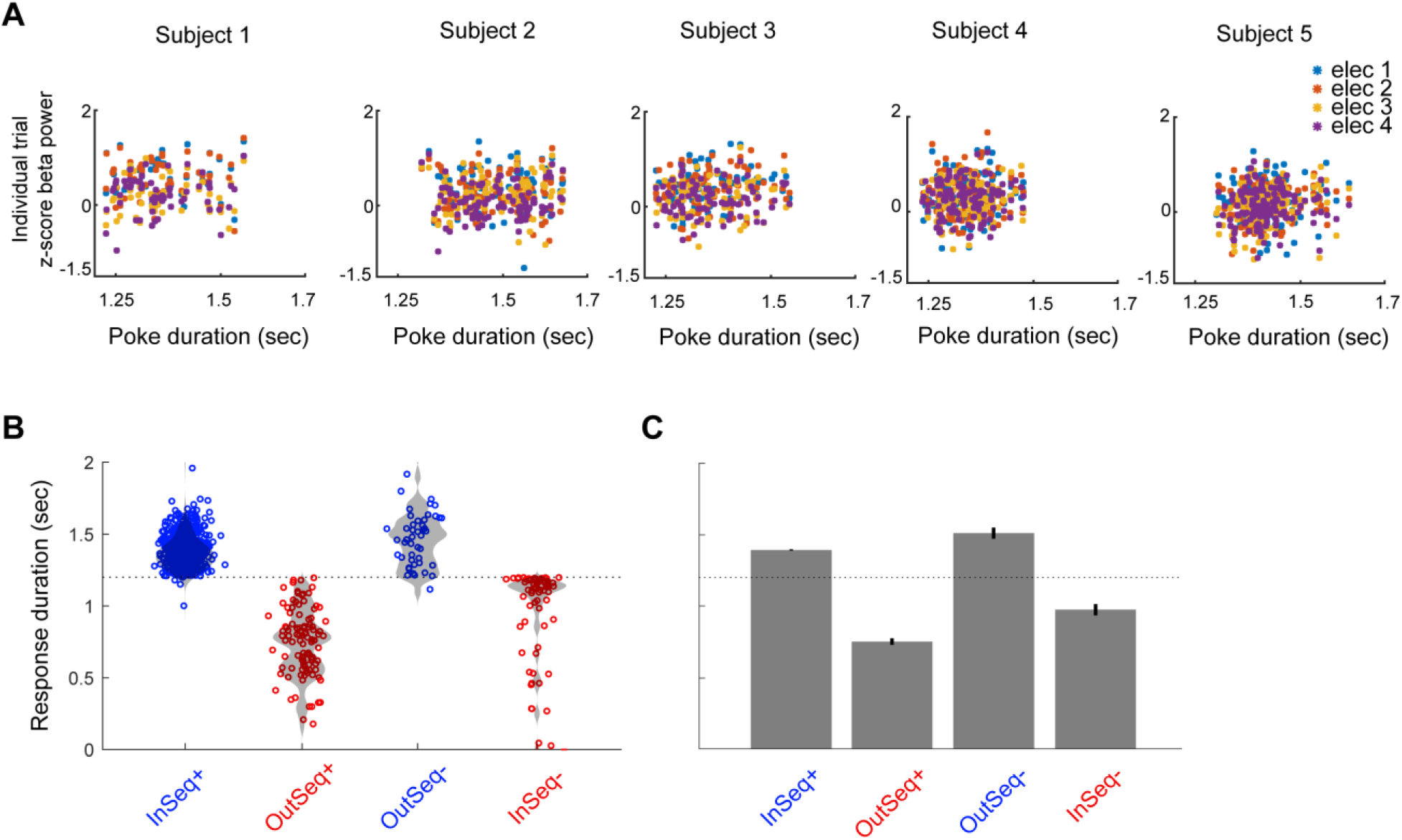
Correlations between 20-40 Hz power and poke duration across trials. **A.** Correlation between 20-40 Hz power (250ms period before port withdrawal) and poke duration across trials (InSeq+ trials only), calculated for each electrode separately. Data from each animal is shown in columns, and data from each animals’ four electrodes is correspondingly color-coded. Correlations centered around 0 were observed, with only 1 out of 20 electrodes showing a significant positive correlation (r = 0.2722, *p* = 0.0008). **B.** Distribution of response durations across trial types, aggregating data across animals. **C.** Mean response durations across trial types from data shown in B.

**Figure S2.**
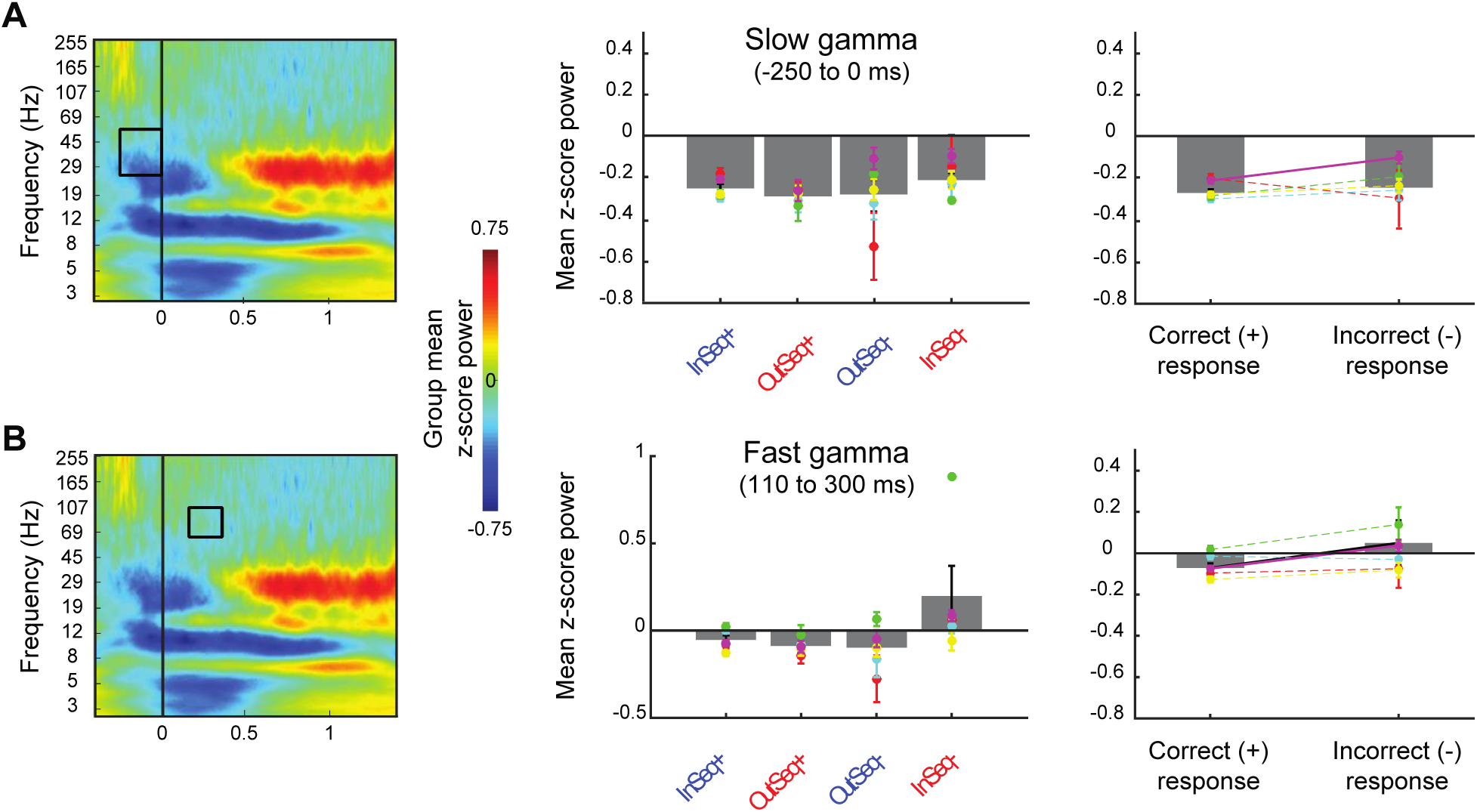
Slow gamma (25-55 Hz) and fast gamma (60-100 Hz) power across trial types. Peri-event group spectrograms relative to port entry during the well-trained session (showing most proximal CA1 site). Time-frequency ranges used for bar plots in second and third columns are indicated with black boxes on the spectrograms in the first column (data averaged across the four electrodes). **A.** Slow gamma power (25-55 Hz) for the 250 ms period preceding port entry (“retrieval” period) across all four trial types (InSeq+, OutSeq-, OutSeq+, and InSeq-) and collapsing across correct (InSeq+ and OutSeq+) and incorrect (InSeq- and OutSeq-) responses. **B.** Fast gamma power (60-100 Hz) for the 110-300 ms time window (“encoding” period) across the same trial types and correct vs incorrect contrast. None of the group-level comparisons were significant (all *p*’s < 0.05) and only one animal showed a significant increase on incorrect trials, which was driven by high values on InSeq-trials (*p* < 0.0001). See Table S5 for permutation testing statistical report.

## Supplementary Tables

**Table S1.** Odor Processing Electrode-Pair Permutation Testing and one-way ANOVA Results.

| A. Odor processing 20-40 Hz power, 110-1200 msec (corresponds to Figure 2C) |  |  |  |  |  |  |  |  |
| --- | --- | --- | --- | --- | --- | --- | --- | --- |
| Animal | 1vs2 | 1vs3 | 1vs4 | 2vs3 | 2vs4 | 3vs4 | p(FDR) | ANOVA<br>df=3 |
| 1 | 0.9471 | 0.0020 | 0.0020 | 0.0020 | 0.0020 | 0.9311 | 0.002 | F=53.47<br>p=1.3466e-26 |
| 2 | 0.4795 | 0.0020 | 0.0020 | 0.0020 | 0.0020 | 0.3437 | 0.002 | F=15.15<br>p=2.2445e-09 |
| 3 | 0.8332 | 0.0080 | 0.0080 | 0.0040 | 0.0040 | 0.8831 | 0.008 | F=5.56<br>p=9.7137e-04 |
| 4 | 0.7852 | 0.2657 | 0.6334 | 0.4316 | 0.9131 | 0.5195 | 0.000 | F=0.42<br>p=0.7395 |
| 5 | 0.2537 | 0.0040 | 0.0080 | 0.0200 | 0.0579 | 0.6893 | 0.020 | F=4.97<br>p=0.0020 |
| B. Odor processing 4-8 Hz power, 110-1200 msec (corresponds to Figure 2D) |  |  |  |  |  |  |  |  |
| Animal | 1vs2 | 1vs3 | 1vs4 | 2vs3 | 2vs4 | 3vs4 | p(FDR) | ANOVA<br>df=3 |
| 1 | 0.5375 | 0.0020 | 0.0020 | 0.0040 | 0.0020 | 0.6214 | 0.0040 | F=11.17<br>p=6.7785e-07 |
| 2 | 0.0020 | 0.8651 | 0.5235 | 0.000 | 0.000 | 0.6553 | 0.0020 | F=12.14<br>p=1.2516e-07 |
| 3 | 0.0040 | 0.000 | 0.0340 | 0.000 | 0.000 | 0.0080 | 0.0340 | F=23.03<br>p=1.1269e-13 |
| 4 | 0.000 | 0.000 | 0.000 | 0.000 | 0.000 | 0.9051 | 0.0000 | F=26.79<br>p=3.8771e-16 |
| 5 | 0.000 | 0.000 | 0.000 | 0.000 | 0.040 | 0.0719 | 0.0400 | F=119.8<br>p=9.9337e-62 |
| **Note:** (Columns 2-7) Electrode pairwise permutation test p-values; (Column 8) Thresholded p-value for permutation test FDR correction for multiple comparisons; (Column 9) Electrode-20-40hz power main effect (ANOVA with reported F-statistic (df=3) and p-value. |  |  |  |  |  |  |  |  |

**Table S2.** Running Electrode-Pair Permutation Testing and one-way ANOVA Results.

| A. Running >24 Hz power (corresponds to Figure 2E) |  |  |  |  |  |  |  |  |
| --- | --- | --- | --- | --- | --- | --- | --- | --- |
| Animal | 1vs2 | 1vs3 | 1vs4 | 2vs3 | 2vs4 | 3vs4 | p(FDR) | ANOVA<br>df=3 |
| 1 | 0.8711 | 0.0200 | 0.0779 | 0.0100 | 0.0639 | 0.0000 | 0.020 | F=6.49<br>p=2.5445e-04 |
| 2 | 0.3277 | 0.0000 | 0.0000 | 0.0000 | 0.0000 | 0.1099 | 0.0000 | F=53.63<br>p=5.6225e-30 |
| 3 | 0.0280 | 0.0000 | 0.0000 | 0.0000 | 0.0000 | 0.0000 | 0.0280 | F=270.23<br>p=1.1893e-97 |
| 4 | 0.0799 | 0.0000 | 0.0000 | 0.0000 | 0.0000 | 0.0639 | 0.0000 | F=34.19<br>p=3.1060e-20 |
| 5 | 0.3117 | 0.4476 | 0.0000 | 0.6613 | 0.0000 | 0.0000 | 0.0000 | F=12.46<br>p=6.2672e-08 |
| B. Running 4-12 Hz power (corresponds to Figure 2F) |  |  |  |  |  |  |  |  |
| Animal | 1vs2 | 1vs3 | 1vs4 | 2vs3 | 2vs4 | 3vs4 | p(FDR) | ANOVA<br>df=3 |
| 1 | 0.000 | 0.2557 | 0.0739 | 0.0020 | 0.0020 | 0.4076 | 0.0020 | F=16.3<br>p=3.7733e-10 |
| 2 | 0.5594 | 0.1998 | 0.4535 | 0.5894 | 0.2058 | 0.0639 | 0.0000 | F=1.33<br>p=0.2650 |
| 3 | 0.0020 | 0.0020 | 0.0020 | 0.0659 | 0.0020 | 0.0480 | 0.0020 | F=21.91<br>p=3.8251e-13 |
| 4 | 0.0020 | 0.0020 | 0.0020 | 0.0519 | 0.0020 | 0.0020 | 0.0020 | F=54.53p=8.97<br>91e-31 |
| 5 | 0.0679 | 0.0260 | 0.0659 | 0.6613 | 0.8911 | 0.7952 | 0.0000 | F=2.14<br>p=0.0940 |
| **Note:** (Columns 2-7) Electrode pairwise permutation test p-values; (Column 8) Thresholded p-value for permutation test FDR correction for multiple comparisons; (Column 9) Electrode-20-40hz power main effect (ANOVA with reported F-statistic (df=3) and p-value. |  |  |  |  |  |  |  |  |

**Table S3.** 20-40 Hz Power Variation Along Proximodistal Axis and with Sequence Knowledge.

| A. 20-40 Hz power along the proximodistal axis (corresponds to Figure 3B) |  |  |  |  |  |  |  |  |
| --- | --- | --- | --- | --- | --- | --- | --- | --- |
| Animal | 1vs2 | 1vs3 | 1vs4 | 2vs3 | 2vs4 | 3vs4 | p(FDR) | ANOVA<br>df=3 |
| 1 | 0.8531 | 0.002 | 0.0020 | 0.002 | 0.002 | 0.9431 | 0.0020 | F=20.08<br>P=9.5650e-12 |
| 2 | 0.1499 | 0.0380 | 0.0020 | 0.6933 | 0.0040 | 0.0060 | 0.0060 | F=8.28<br>P=2.28299e-05 |
| 3 | 0.4915 | 0.0020 | 0.0020 | 0.0020 | 0.0020 | 0.2158 | 0.0020 | F=14.26<br>P=8.2435e-09 |
| 4 | 0.9231 | 0.3357 | 0.2897 | 0.3017 | 0.2378 | 0.9970 | 0.0000 | F=0.76<br>P=0.5159 |
| 5 | 0.1938 | 0.8152 | 0.8871 | 0.6374 | 0.2997 | 0.7173 | 0.0000 | F=5.52<br>P=0.001 |
| B. 20-40 hz power with sequence knowledge (corresponds to Figure 3C) |  |  |  |  |  |  |  |  |
| Animal | Novel 1<br>vs. Novel 2 |  | Well-trained vs.<br>Novel 1 |  | Well-trained vs.<br>Novel2 |  | p(FDR) | ANOVA<br>df=3 |
| 1 | 0.007992 |  | 0.0000 |  | 0.001998 |  | 0.0080 | F=15.79<br>P=3.53472e-07 |
| 2 | 0.37962 |  | 0.03996 |  | 0.13187 |  | 0.0000 | F=2.55<br>P=0.08 |
| 3 | 0.33367 |  | 0.063936 |  | 0.32168 |  | 0.0000 | F=1.77<br>P=0.1727 |
| 4 | 0.017982 |  | 0.11788 |  | 0.001998 |  | 0.0180 | F=10.74<br>P=2.9866e-05 |
| **Note:** (Columns 2-7) Electrode pairwise permutation test p-values; (Column 8) Thresholded p-value for permutation test FDR correction for multiple comparisons; (Column 9) Electrode-20-40hz power main effect (ANOVA with reported F-statistic (df=3) and p-value. |  |  |  |  |  |  |  |  |

**Table S4.** Trial Count for Individual Animals.

| Animal | InSeq+ | OutSeq+ | InSeq- | OutSeq- |
| --- | --- | --- | --- | --- |
| 1 | 73 | 24 | 3 | 3 |
| 2 | 114 | 22 | 12 | 5 |
| 3 | 101 | 11 | 1 | 10 |
| 4 | 132 | 27 | 13 | 15 |
| 5 | 164 | 16 | 25 | 14 |
| TOTAL | 584 | 100 | 54 | 47 |

**Table S5.** P-values (permutation test) for power means in different frequency ranges across trial types.

| Correct vs. incorrect | Anim1 | Anim2 | Anim3 | Anim4 | Anim5 |
| --- | --- | --- | --- | --- | --- |
| 20-40 Hz (beta) | 0.3237 | 0.9730 | 0.5055 | 0.2258 | 0.7592 |
| 25-55 Hz (slow gamma) | 0.3736 | 0.4136 | 0.1798 | 0.2098 | 0.0000 |
| 60-100 Hz (fast gamma) | 0.8092 | 0.7133 | 0.0619 | 0.2218 | 0.0000 |
| 4-8 Hz (theta) | 0.8931 | 0.7213 | 0.3576 | 0.3816 | 0.1838 |

## References

Agster KL, Burwell RD (2009) Cortical efferents of the perirhinal, postrhinal, and entorhinal cortices of the rat. Hippocampus 19(12):1159–86.

Allen TA, Fortin NJ (2013) The evolution of episodic memory. Proc Natl Acad Sci U S A 110:10379 – 10386.

Allen TA, Salz DM, McKenzie S, Fortin NJ (2016) Nonspatial sequence coding in CA1 neurons. J Neurosci 36(5):1547–1563.

Buzsáki G, Logothetis N, Singer W (2013) Scaling brain size, keeping timing: evolutionary preservation of brain rhythms. Neuron 80(3):751–64.

Buzsáki G, Tingley D (2018) Space and Time: The Hippocampus as a Sequence Generator. Trends Cogn Sci 22(10):853–869.

Colgin LL (2016) Rhythms of the hippocampal network. Nature Rev Neuro 17(4):239–249.

Colgin LL, Denninger T, Fyhn M, Hafting T, Bonnevie T, Jensen O, Moser MB, Moser EI (2009) Frequency of gamma oscillations routes flow of information in the hippocampus. Nature 462(7271):353–7.

Eichenbaum H (2014) Time cells in the hippocampus: a new dimension for mapping memories. Nat Rev Neurosci 15:732–744.

Engel AK, Fries P (2010) Beta-band oscillations--signalling the status quo? Curr Opin Neurobiol 20(2):156–65.

Fortin NJ, Agster KL, Eichenbaum HB (2002) Critical role of the hippocampus in memory for sequences of events. Nat Neurosci 5(5):458–62.

Foster DJ, Knierim JJ (2012) Sequence learning and the role of the hippocampus in rodent navigation. Curr Opin Neurobiol 22(2):294–300.

Haberly LB, Price JL (1978) Association and commissural fiber systems of the olfactory cortex of the rat. J Comp Neurol 178(4):711–40.

Hartzell AL, Burke SN, Hoang LT, Lister JP, Rodriguez CN, Barnes CA (2013) Transcription of the immediate-early gene Arc in CA1 of the hippocampus reveals activity differences along the proximodistal axis that are attenuated by advanced age. J Neurosci 33 (8), 3424–3433.

Hasselmo ME, Wyble BP (1997) Free recall and recognition in a network model of the hippocampus: simulating effects of scopolamine on human memory function. Behav Brain Res 89:1–34.

Henriksen EJ, Colgin LL, Barnes CA, Witter MP, Moser MB, Moser EI (2010) Spatial representation along the proximodistal axis of CA1. Neuron 68 (1) (2010) 127–137.

Igarashi KM, Lu L, Colgin LL, Moser MB, Moser EI (2014) Coordination of entorhinal-hippocampal ensemble activity during associative learning. Nature 510:143–147.

Kesner RP, Gilbert PE, Barua LA (2002) The role of the hippocampus in memory for the temporal order of a sequence of odors. Behav Neurosci 116(2):286–90.

Kilavik BE, Zaepffel M, Brovelli A, MacKay WA, Riehle A (2013) The ups and downs of beta oscillations in sensorimotor cortex. Exp Neurol 245: 15–26.

Lisman JE (1999) Relating hippocampal circuitry to function: recall of memory sequences by reciprocal dentate-CA3 interactions. Neuron 22(2):233–42.

Lisman JE, Grace AA (2005) The hippocampal-VTA loop: controlling the entry of information into long-term memory. Neuron 46:703–713.

MacDonald CJ, Lepage KQ, Eden UT, Eichenbaum H (2011) Hippocampal “time cells” bridge the gap in memory for discontiguous events. Neuron 71:737–749.

Martin C, Beshel J, Kay LM (2007) An Olfacto-Hippocampal Network Is Dynamically Involved in Odor-Discrimination Learning. J Neurophys 98(4):2196–2205.

Martin C and Ravel N (2014). Beta and gamma oscillatory activities associated with olfactory memory tasks: different rhythms for different functional networks? Front Behav Neurosci 8, 218

McNaughton BL, Morris RGM (1987) Hippocampal synaptic enhancement and information storage within a distributed memory system. Trends Neurosci 10:408–415.

Miller EK, Lundqvist M, Bastos AM (2018) Working memory 2.0. Neuron 100: 463–475.

Nakazawa Y, Pevzner A, Tanaka KZ, Wiltgen BJ (2016) Memory retrieval along the proximodistal axis of CA1. Hippocampus 26 (9), 1140–1148.

Ng CW, Elias GA, Asem JSA, Allen TA, Fortin NJ (2018) Nonspatial sequence coding varies along the CA1 transverse axis. Behav Brain Res: 354:39–47.

O’Keefe J, Nadel L (1978) The Hippocampus as a Cognitive Map. Clarendon Press: Oxford.

Pesaran B, Vinck M, Einevoll GT, Sirota A, Fries P, Siegel M, Truccolo W, Schroeder CE, Srinivasan R (2018) Investigating large-scale brain dynamics using field potential recordings: analysis and interpretation. Nat Neurosci 21:1–17.

Robinson NTM, Priestley JB, Rueckemann JW, Garcia AD, Smeglin VA, Marino FA, Eichenbaum HB (2017) Medial entorhinal cortex selectively supports temporal coding by hippocampal neurons. Neuron 94(3): 677–688.e6.

Schmidt R, Ruiz MH, Kilavik BE, Lundqvist M, Starr PA, Aron AR (2019) Beta oscillations in working memory, executive control of movement and thought and sensorimotor function. J Neurosci 39(42), 8231–8238.

Shahbaba B, Li L, Agostinelli F, Saraf M, Elias GA, Baldi P, Fortin NJ (2019) Hippocampal ensembles represent sequential relationships among discrete nonspatial events. bioRxiv https://doi.org/10.1101/840199

van Strien NM, Cappaert NL, Witter MP (2009) The anatomy of memory: an interactive overview of the parahippocampal-hippocampal network. Nat Rev Neurosci 10(4):272–82.

Wiener M, Parikh H, Krakow A, Coslett HB (2018) An intrinsic role of beta oscillations in memory for time estimation. Scientific Reports 8(1), 7992.

Witter MP, Doan TP, Jacobsen B, Nilssen ES, Ohara S (2017) Architecture of the Entorhinal Cortex A Review of Entorhinal Anatomy in Rodents with Some Comparative Notes. Front Syst Neurosci 11:46.

